# Oxidative stress underlies heritable impacts of paternal cigarette smoke exposure

**DOI:** 10.1101/750638

**Authors:** Patrick J Murphy, Jingtao Guo, Timothy G Jenkins, Emma R James, John R Hoidal, Thomas Huecksteadt, Dallin Broberg, James M Hotaling, David F Alonso, Douglas T Carrell, Bradley R Cairns, Kenneth I Aston

## Abstract

Paternal cigarette smoke (CS) exposure is associated with increased risk of behavioral disorders and cancer in offspring, but the mechanism has not been identified. This study used mouse models to evaluate: 1) what impact paternal CS exposure has on sperm DNA methylation (DNAme), 2) whether sperm DNAme changes persist after CS exposure ends, 3) the degree to which DNAme and gene expression changes occur in offspring and 4) the mechanism underlying impacts of CS exposure. We demonstrate that CS exposure induces sperm DNAme changes that are partially corrected within 28 days of removal from CS exposure. Additionally, paternal smoking causes changes in neural DNAme and gene expression in offspring. Remarkably, the effects of CS exposure are largely recapitulated in oxidative stress-compromised *Nrf2*^-/-^ mice and their offspring, independent of paternal smoking. These results demonstrate that paternal CS exposure impacts offspring phenotype and that oxidative stress underlies CS induced heritable epigenetic changes.

## INTRODUCTION

Cigarette smoke (CS) exposure is a global epidemic with significant health consequences. In a recent study it was estimated that more than one third of the world’s population is regularly exposed to tobacco smoke (Oberg et al., 2011). The health consequences of smoke exposure are significant and include numerous diseases and dysfunctions of the respiratory tract, increased risk of multiple types of cancer and cardiovascular disease (DiFranza et al., 2004, Moritsugu, 2007). Tobacco smoke exposure is a global problem, the implications of which are becoming increasingly apparent. However, little is known about the impact of paternal exposure to CS on sperm and implications of pre-conception paternal CS on offspring health (Laubenthal et al., 2012).

Male fertility rates have steadily declined in developed countries over the past half-century (Rolland et al., 2012, Priskorn et al., 2012, Swan et al., 1997). These trends are due to a variety of factors, but increased exposures to environmental toxins and negative lifestyle factors likely contribute (Kiziler et al., 2007, Kulikauskas et al., 1985). Cigarette smoking is associated with an accumulation of cadmium and lead in seminal plasma, reduced sperm count and motility, and increased morphological abnormalities in sperm (Kiziler et al., 2007, Kulikauskas et al., 1985). In addition, reduced reproductive potential has been reported in tobacco smoke-exposed mice (Polyzos et al., 2009) and humans (Fuentes et al., 2010). Adult male mice exposed to sidestream tobacco smoke display significant increases in sperm DNA mutations at expanded simple tandem repeats (ESTRs) (Marchetti et al., 2011), as well as more frequent aberrations in sperm chromatin structure and elevated sperm DNA damage (Polyzos et al., 2009). In contrast, CS-exposed male mice exhibit no measurable increase in somatic cell chromosome damage, indicating that germ cells may be more prone to environmentally-induced genetic and/or epigenetic insults compared with somatic cells.

The International Association for Research on Cancer has declared that *paternal* smoking *prior* to pregnancy is associated with a significantly elevated risk of leukemia in the offspring (Secretan et al., 2009), suggesting CS-induced changes occur in sperm that influence offspring phenotype. Smoking has been clearly shown to modify DNAme patterns and gene expression in somatic tissues in individuals exposed to first- or second-hand tobacco smoke (Kohli et al., 2012, Bosse et al., 2012, Word et al., 2012) as well as in newborns of smoking mothers (Breton et al., 2009, Perera and Herbstman, 2011, Joubert et al., 2012). We recently reported altered sperm DNA methylation (DNAme) patterns in men who smoke (Jenkins et al., 2017).

Mounting evidence supports sperm epigenetic changes as a mechanism for increased health risks in offspring of smoking fathers. We therefore aimed to explore the dynamics of sperm epigenetic changes after withdrawal from smoke, and to explore the mechanism underlying CS-induced alterations in offspring.

## RESULTS

### Experimental design and phenotypic effects

Male mice were assigned to CS-exposed or non-exposed groups (n = 10-12 per group). The CS animals were exposed to the body mass-adjusted equivalent of 10-20 cigarettes per day, 5 days per week over a period of 60 days-corresponding to two complete cycles of spermatogenesis (Figure 1A). CS-exposed and control mice were bred to unexposed females, and offspring were analyzed for phenotypic and molecular measures. The nuclear factor (erythroid-derived 2)-like 2 (NRF2) pathway is the primary cellular defense against the cytotoxic effects of oxidative stress. Thus, to investigate the mode by which CS-induced epigenetic changes occur, we utilized the *Nrf2*^*-/-*^ mouse model, which has compromised antioxidant capacity. In agreement with the literature, both wild type (WT) and *Nrf2*^*-/-*^ mice that were exposed to CS weighed significantly less than non-exposed control animals (Figure S1A). Sperm concentration and motility were not significantly impacted by CS exposure, but sperm concentration was lower in *Nrf2*^*-/-*^ than WT males, and conception was significantly delayed in CS-exposed *Nrf2*^*-/-*^mice compared with unexposed *Nrf2*^*-/-*^mice (Figure S1A). Neither growth trajectories nor sperm parameters were different in F1 animals based on paternal CS exposure (Figure S1B).

**Figure 1.**
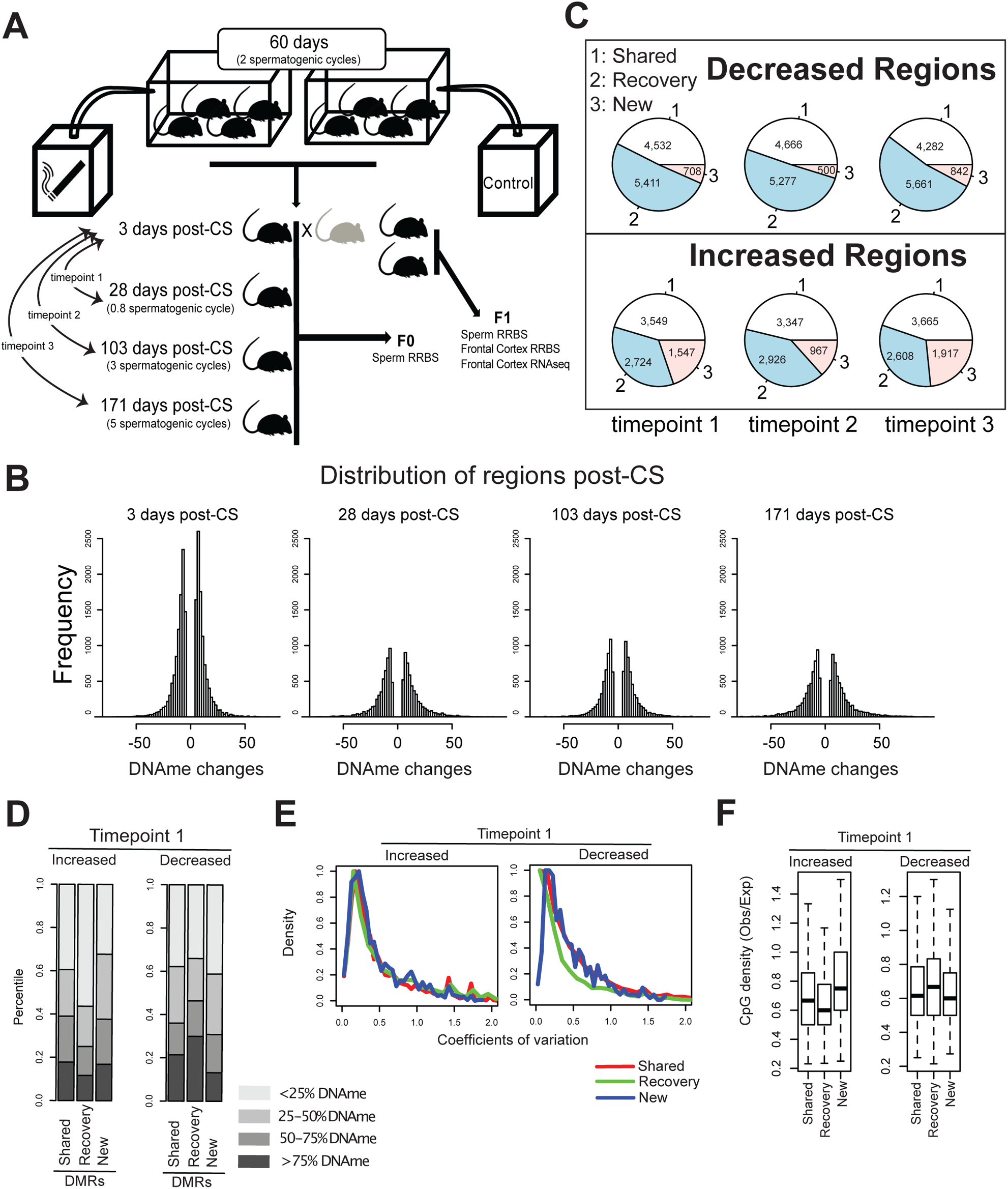
Schematic of study design and regional DNA methylation changes and recovery. **A)** Male mice were assigned to CS-exposed or non-exposed groups (n = 10-12 per group). The CS animals were exposed to the body mass-adjusted equivalent of 10-20 cigarettes per day, 5 days per week over a period of 60 days-corresponding to two complete cycles of spermatogenesis. CS-exposed and control mice were bred to unexposed females, and offspring were analyzed for phenotypic and molecular measures (see method for more details). **B)** Histograms describing the changes of DNAme in sperm after CS-exposure and recovery. **C)** The majority of DMRs observed prior to recovery were either maintained across the recovery period or returned to baseline levels, with only a small fraction of new DMRs emerging during the recovery period. Comparison is based on the 3 days post-CS group. **D)** Impact of the initial methylation status and direction of change on methylation recovery in the group analyzed 28 days after removal of CS. The hypomethylated DMRs (<25% DNAme) in which methylation increased and the hypermethylated DMRs (> 75% DNAme) in which methylation decreased with smoke exposure were more likely to recover. Regions of intermediate DNAme were less likely to recover. **E)** DMR recovery as a function of CpG density in the 28 days post-CS group. In every category of DMR (shared, recovery or new) variation diminished as CpG density of a region increased. The impact of CpG density on variation was particularly apparent for regions in which DNAme decreased as a result of CS exposure and later recovered to baseline. **F)** DMRs that displayed increased DNAme in CS-exposed animals and recovered within 28 days post-CS were generally regions of lower CpG density, while DMRs that lost methylation and subsequently recovered were generally at regions of higher CpG density.

### Smoking-induced DNAme changes in F0 sperm recovered in accordance with CG density

We performed reduced representation bisulfite sequencing (RRBS) to explore the effects of smoking on F0 sperm DNAme (sperm collected within 3 days of completing a 60-day smoking treatment), and we examined whether CS-induced DNAme changes recover to baseline unexposed levels following removal of CS exposure (28, 103 and 171 days after smoking treatment; n = 10 per group). Due to the epigenetic reprogramming that occurs in early embryos, it is unlikely that paternal sperm DNAme directly influences adult offspring phenotype. However, because DNAme is a sensitive marker and can be reliably assessed on a genome-wide scale, we view it an important signature of epigenetic change.

We found CS-associated changes in DNAme at a large number of individual CpGs (Figure S2A) with essentially equal representation of sites that lost DNAme and gained DNAme. In addition, we found that the number of differentially methylated CpGs declined only slightly after removal of CS exposure for 28-171 days (Figure S2A). We partitioned differentially methylated CpGs into three groups: 1) shared, meaning differentially methylated CpGs that were maintained between the treatment group and the respective recovery group, 2) recovered, meaning CpGs that were differentially methylated after initial exposure but were no longer differentially methylated in the recovery group, and 3) new, meaning CpGs that were not differentially methylated after initial exposure but emerged as newly differentially methylated in the recovery groups. These classes were assessed separately for CpGs that initially lost DNAme verses those that initially gained DNAme relative to the control. For CpGs that lost DNAme, the three groups were relatively evenly represented across recovery times, with only a slight over-representation of shared CpGs. For CpGs that gained DNAme, there was a slight bias toward the emergence of new differentially methylated CpGs across recovery groups (Figure S2B).

Given that DNAme status of CpG regions likely has a greater capacity to confer functional effects compared with individual sites, we subsequently binned individual CpGs into regions based on their proximity (see methods). We then performed analyses similar to those performed for individual CpGs. As expected, changes in DNAme occurred at far fewer regions compared with individual CpGs (Figure S4A), and importantly, we found strong evidence for recovery of smoke-induced sperm DNAme changes following removal of the exposure. Indeed, the majority of DNAme regions that recovered returned to control levels within 28 days of removal of smoke exposure, and additional recovery was not observed following longer recovery periods (Figure 1B). In contrast with the individual CpG data, we found that far fewer new differentially methylated regions (DMRs) emerged during the recovery period (Figure 1C).

We then sought to understand epigenetic properties that might impact the ability of specific regions to recover following removal of the CS. For this analysis, we classified all DMRs into six classes, based on their dynamics during recovery (shared, recovered or new) and their direction of change (increase or decrease). We discovered that upon smoke exposure the hypomethylated regions where DNAme increased, and hypermethylated regions where DNAme decreased, were more likely to recover (Figures 1D and S2C). Intermediate DNAme were less likely to recover across all groups. In addition, we observed that recovered DMRs that initially decreased in DNAme level show lower variation compared to all other groups (Figures 1E and S2D), thus suggesting that regions of extreme hyper- and hypo-DNAme were less likely to change after CS exposure, and when changes did occur, these regions were more likely to recover. Moreover, DMRs that displayed increased DNAme in CS-exposed animals and subsequently recovered were generally regions of lower CpG density, while DMRs that lost DNAme and subsequently recovered were generally at regions of higher CpG density (Figures 1F and S2E). Our observation that individual CpGs, and low density regions did not recover to nearly the same degree as CpG dense regions suggests that CpG density might function to “buffer” against environmental insults.

### Paternal CS exposure impacted DNAme and gene expression changes in F1 brains

We next sought to determine whether paternal CS exposure had any effect on offspring phenotype. Given that brain and nervous systems were previously reported to be sensitive to preconception paternal exposures (Dai et al., 2017, Hawkey et al., 2019, Pachenari et al., 2019, Levin et al., 2019), we investigated DNAme by RRBS in the prefrontal cortex of F1 mice derived from CS-exposed sires compared to the offspring of unexposed males (n = 8 per group). We found that paternal smoking altered DNAme patterns in over 28,000 regions in the prefrontal cortex of offspring (Figure 2C-D).

**Figure 2.**
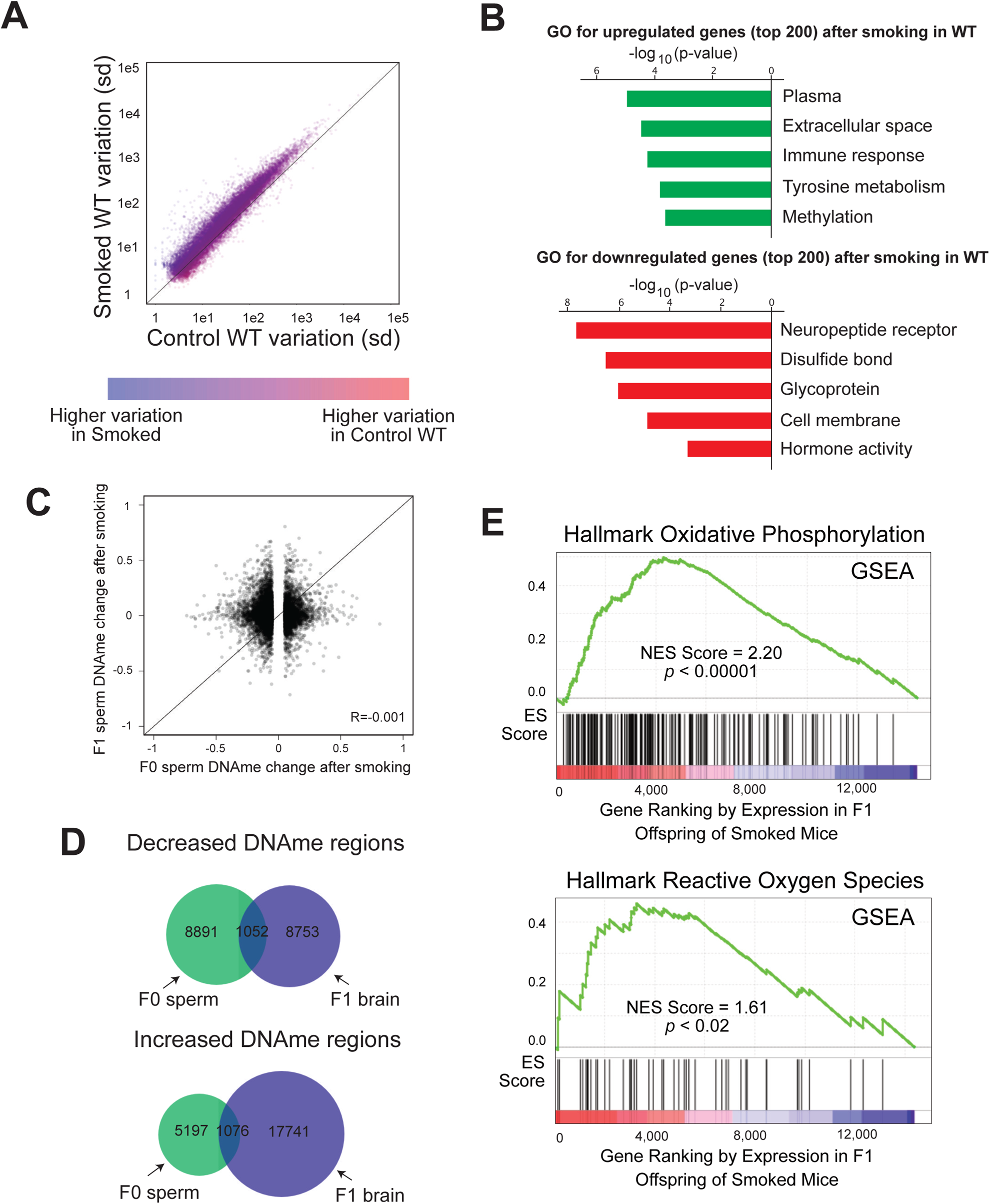
Cigarette smoking leads to changes in gene expression in offspring brain, but these changes do not suggest a direct DNAme inheritance model. **A)** Variation in F1 prefrontal cortex gene expression was significantly higher in CS-exposed animals compared with controls suggesting stochastic dysregulation of gene expression in paternal CS-exposed offspring and offspring of mice with reduced antioxidant capacity. **B)** Gene ontology analysis of significantly upregulated and downregulated genes associated with paternal CS exposure indicated significant overrepresentation of several gene families. **C)** The DNAme changes observed in F0 sperm were not observed in the sperm of F1 offspring, suggesting the CS-associated effects likely do not confer risk beyond the first generation. **D)** Likewise, no significant overlap was observed in DMRs in F0 sperm compared with DMRs in F1 prefrontal cortex. **E)** GSEA analysis revealed that oxidative phosphorylation pathway genes (top) and reactive oxygen species genes (bottom) are significantly upregulated in the brains of the F1 offspring of smoked mice.

To further investigate the potential for phenotypic effects in offspring associated with paternal CS exposure, we performed RNA-seq of the F1 prefrontal cortex in the same animals as those assessed for DNAme. Interestingly, paternal CS exposure caused a globally elevated variation in gene expression in the F1 brains (Figure 2A). This increased variation limited our ability to reliably identify differentially expressed genes. Instead, we ranked genes by changes in gene expression and performed gene set enrichment analysis (GSEA) and gene ontology (GO) analysis. Using GSEA, we found that gene expression increased for genes associated with oxidative stress (Figure 2E). Using GO analysis, gene transcripts associated with gene classes including immune response and metabolism were over-represented, and transcripts associated with neuropeptide receptors and hormone activity were significantly under-represented in the offspring of CS-exposed males (Figure 2B).

### Paternal CS-induced effects are likely not transmitted beyond the first generation

To investigate whether the impact of CS exposure could be passed to additional subsequent generations, we compared the smoke-associated DNAme changes in F0 sperm with those of F1 sperm and found no correlation (r=-0.001; Figure 2C). As expected, we also found that paternal CS-associated DMRs in F1 brains and those in smoke-exposed F0 sperm had minimal overlap (hypergeometric p-value =1; Figure 2D), indicating that F0 sperm DNAme changes do not directly impact DNAme levels in the F1 prefrontal cortex, consistent with the DNAme erasure that occurs during early mouse development. Accordingly, regions where DNAme changes occurred are not marked by chromatin features that are known to be maintained in mature sperm (Jung et al., 2017) (Figure S3A-C). While the DNAme effects described here can certainly be considered a marker of epigenetic impacts of CS, these results suggest that the effects of CS exposure likely do not persist beyond the first generation.

### CS-induced epigenetic changes were regulated by oxidative stress

Based on our GSEA studies (Figure 2E), we hypothesized that impacts of CS exposure might be due in part to oxidative stress. This led us to examine F0 sperm DNAme changes in *Nrf2*^*-/-*^ mice (compared to WT mice), as well as CS-exposed mice compared with unexposed *Nrf2*^*-/-*^ mice. If oxidative stress played a role in the sperm DNAme changes, the changes observed in *Nrf2*^*-/-*^ mice would mirror changes in CS-exposed WT mice, and impact on CS-exposed *Nrf2*^*-/-*^ mice might be more pronounced. In agreement with our hypothesis, we found that CS-induced DNAme changes in WT sperm were largely recapitulated in *Nrf2*^*-/-*^ sperm, but CS exposure seemed to have no additional impact on *Nrf2*^*-/-*^ mouse sperm (Figure 3A). Notably, we did not observe this correlation when comparing our data to similar datasets evaluating the effect of other environmental factors (vinclozolin exposure and protein restricted diet) on sperm DNAme, suggesting independent mechanisms function during CS-exposure (Holland et al., 2016, Brieno-Enriquez et al., 2015) (Figure S3D).

**Figure 3.**
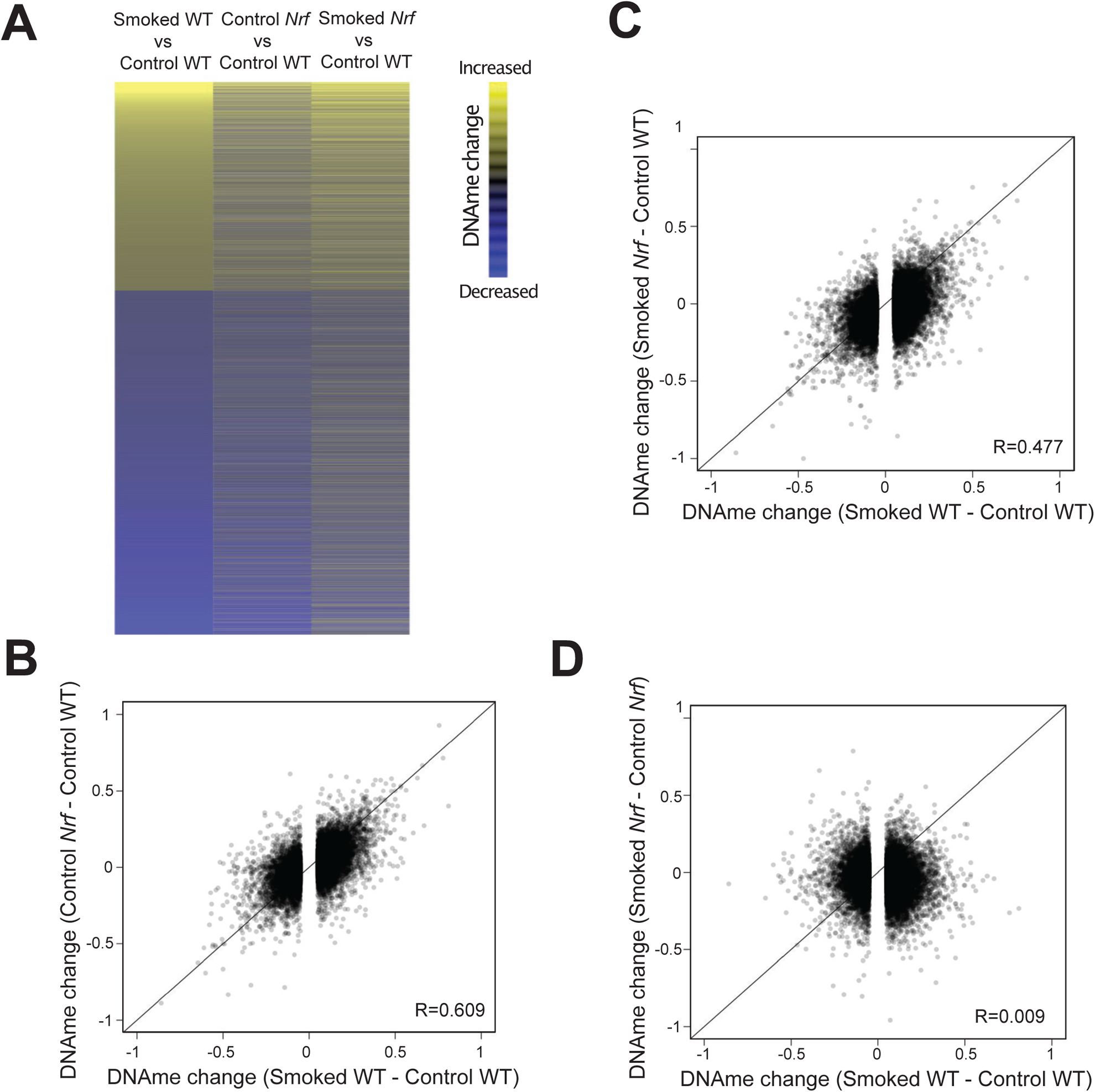
Differential methylation in F0 sperm and F1 prefrontal cortex. **A)** Heatmap illustrating the significant similarity between CS-associated sperm DMRs identified in WT mice and DMRs associated with the *Nrf2*^-/-^ genotype, apparently independent of CS-exposure status. **B)** A highly significant correlation was observed in F1 brain DNAme changes induced by CS exposure in WT sires (x-axis) compared with DNAme changes associated with paternal *Nrf2* status, even in the absence of CS-exposure. **C)** A similar correlation in DNAme change was observed in the offspring of CS-exposed *Nrf2*^+*/-*^mice. **D)** The correlation disappeared when evaluating differential methylation in *Nrf2*^+*/-*^ offspring based on CS exposure status compared with CS-associated DNAme changes in offspring sired by WT mice.

### The oxidative stress effect in *Nrf2*^*-/-*^ offspring closely mirrors changes observed in offspring of CS-exposed mice

Based on our prior results that CS-exposure impacts DNAme and gene expression patterns of offspring, we wondered whether these CS-induced F1 effects might be similar to impacts observed in offspring of *Nrf2*^*-/-*^ males. Indeed, the DNAme changes we observed in *Nrf2*^*-/-*^ F1 animals were highly similar to changes observed in the offspring of CS-exposed WT mice (r=0.609 and r=0.477; Figures 3B and 3C respectively). In agreement with F0 sperm DNAme data, exposure to CS appeared to have no additional impact on brain DNAme beyond the *Nrf2*^*-/-*^ effect (Figure 3D). These observations strongly suggest that CS-induced DNAme changes that occur in F1 brains are mediated by paternal oxidative stress (Venugopal and Jaiswal, 1998, Ishii et al., 2000).

We next sought to determine whether the CS-induced impact on gene expression patterns in F1 brain could also be a consequence of oxidative stress in F0 animals. Similar to our observations in the offspring of CS-exposed WT males, we found globally elevated variation in prefrontal cortex gene expression in the *Nrf2*^+*/-*^ offspring (Figure 4A). In addition, we observed a significant correlation in gene expression between offspring of CS-exposed WT mice and control *Nrf2*^*-/-*^ mice (Figure 4B) as well as between CS-exposed WT and *Nrf2*^+*/-*^ offspring (Figure 4C). Similar to our DNAme measurments, the effects were not further elevated in the offspring of smoke-exposed Nrf2^-/-^ mice (Figure 4D). Likewise, the variation in gene expression was elevated in the offspring of CS-exposed WT mice as well as control *Nrf2*^*-/-*^ mice, but CS exposure did not further elevate the increased gene expression variation in *Nrf*^+*/-*^ offspring beyond the effects of associated with genotype alone (Figure S3E), while variation in gene expression was very similar between CS-exposed WT offspring and offspring of unexposed *Nrf*^*-/-*^ mice (Figure S3F). Although the magnitude of gene expression change was modest, the offspring of *Nrf*^*-/-*^ mice showed significant similarities with gene expression changes observed in the offspring of CS-exposed WT mice based on GO analysis (Figure 4E) and highly significant overlap in differentially expressed genes (Figures 4F). Additionally, GSEA analysis indicated gene sets commonly activated in response to inflammation, and associated with cancer were elevated in both sets of F1 offspring (Figure 4G).

**Figure 4.**
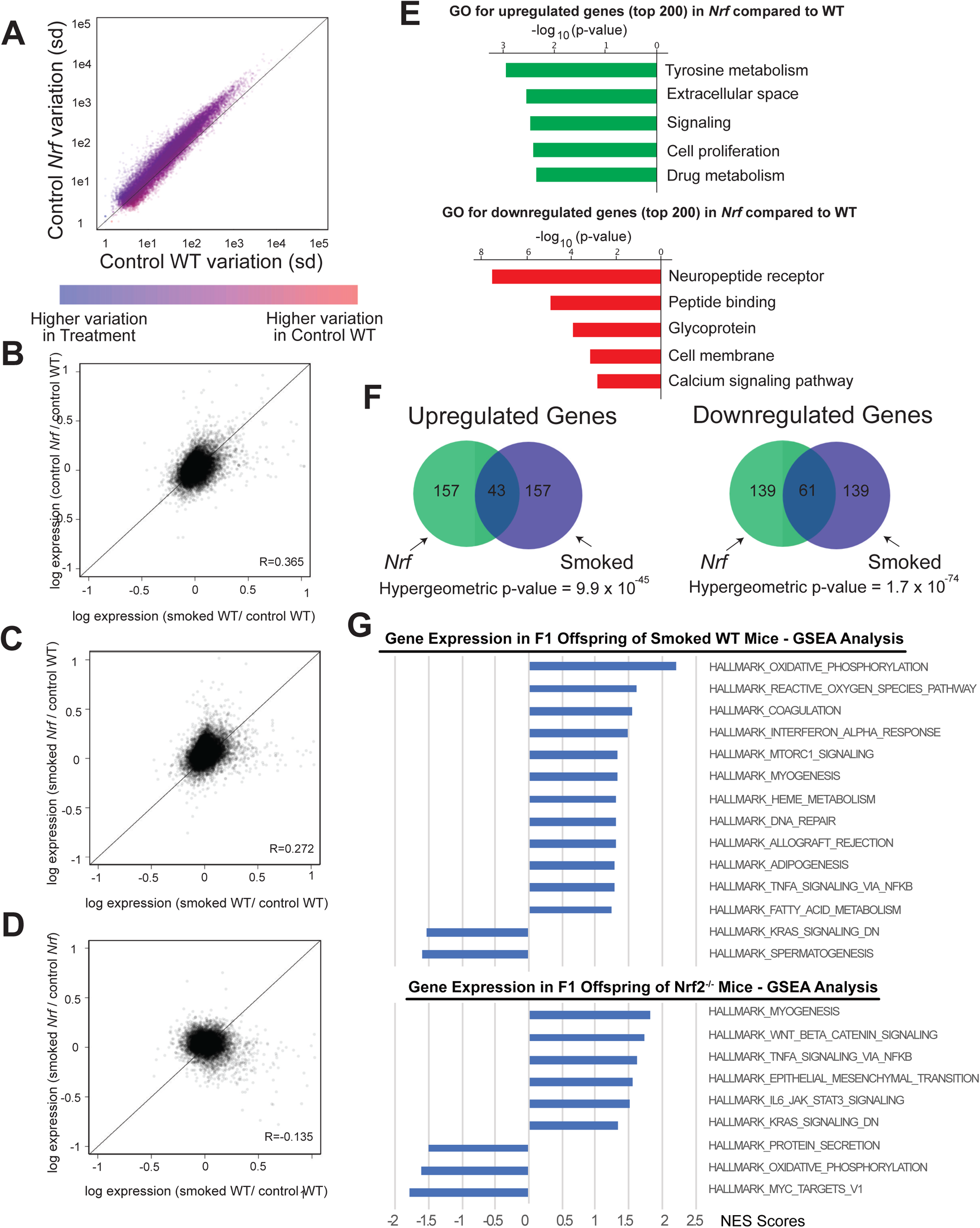
Prefrontal cortex gene expression variation and correlation closely reflect the themes observed in F0 sperm. **A)** Variation in F1 prefrontal cortex gene expression was significantly elevated in *Nrf2*^*-/-*^ controls compared with WT controls suggesting stochastic dysregulation of gene expression in offspring of mice with reduced antioxidant capacity. **B)** A significant correlation in gene expression was observed between offspring of CS-exposed WT mice and control *Nrf2*^*-/-*^ mice. **C)** Likewise, the correlation was observed between CS-exposed WT and *Nrf2*^*-/-*^ offspring. **D)** However, there was no correlation in gene expression changes associated when comparing offspring of CS-exposed and unexposed *Nrf2*^*-/-*^ mice. **E)** GO-terms for genes whose expression change was associated with *Nrf2*^*-/-*^ genotype were similar to those associated with paternal smoking status in WT offspring. **F)** We found a highly significant overlap in differentially expressed genes associated with paternal smoking status in WT animals compared with offspring of unexposed *Nrf2*^*-/-*^ mice. **G)** Pathways that are up- or down-regulated in the offspring brain of smoked WT mice (top) or *Nrf2*^*-/-*^ mice (bottom), based on GSEA analysis.

## DISCUSSION

In this work, we studied the the effect of CS exposure on sperm DNAme. We found many of the DNAme effects that recovered occurred at high CpG dense regions, which are encirhed for genomic regulatory regions (such as promoters and enhancers). Thus, DNAme changes at these regions, which might otherwise impact gene regulation, are likely to be relatively short-lived. These data agree with previous reports that CS exposure significantly impacts DNAme patterns in whole blood, and CS-associated DNAme changes are largely corrected following smoking cessation in a time-dependent manner (Tsaprouni et al., 2014, Zeilinger et al., 2013). We observed recovery after just twenty-eight days, which corresponds to just less than the period of a full spermatogenic cycle in mice of 30 days. In humans, a spermatogenic cycle is 67 days, and additional research is required to characterize the similarities and differences in the dynamics of sperm DNAme alterations between mice and men. Additionally, it is not yet known whether the corrections observed following removal of CS exposure ameliorate the affects observed in offspring, or whether those effects are driven by the regions that persist following CS removal, prompting further investigation.

We further investigated the impact of CS exposure in the oxidative stress-compromised *Nrf2*^-/-^ mice, and found the CS-effects observed in WT animals were largely recapitulated in *Nrf2*^-/-^ mice independent of CS exposure, suggesting elevated oxidative stress as the primary mechanism for CS-mediated sperm epigenetic changes. These findings were consistent with the observations of offspring prefrontal cortex DNAme and gene expression changes. While CS-exposure may represent an extreme example of an environmental insult that induce oxidative stress, the list of environmentally relevant exposures that impact oxidative stress is extensive. The results presented here indicate that all such exposures could potentially impact the epigenetic status of the paternal germline and thus offspring phenotype. In addition, our work provides strong evidence that *Nrf2*^*-/-*^ mice can serve as an important animal model to study CS-exposure induced effects. Further studies may focus on investigating the molecular pathways underlying NRF-mediated DNAme alterations.

By comparing the DNAme and gene expression patterns in the prefrontal cortex between offspring of male mice exposed to CS with those not exposed, we found strong evidence for an impact of paternal smoking on offspring phenotype. We suggest that paternal exposure to inducers of oxidative stress might contribute to behavioral or developmental impacts in offspring. This is supported by GSEA results that showed significant up-regulation of a number of cancer-associated pathways (e.g. INFα, TNFα, Jak-STAT and Wnt/beta catenin; Figure 4G). Notably however, our data showed little overlap of DNAme changes in the F1 prefrontal cortex with DNAme changes in the F0 sperm. While this was somewhat expected, as DNAme state undergoes dramatic reprogramming during early development and neuronal differentiation, it is important to highlight — direct mitotic inheritance of DNAme state is unlikely to mechanistically contribute to the oxidative stress effects we observed in F1 mice. Furthermore, we propose that DNAme may be more accurately described as a marker of epigenetic inheritance and not a mechanistic driver in transmitting environmental impacts to subsequent generations. Additional studies are necessary to confirm this and to distinguish other epigenetic features as markers or drivers, including chromatin status and small RNAs, which could play a significant role in inheritance. Further, the observation that CS-associated sperm DNAme changes were not identified in the sperm of the F1 generation offers reassuring evidence that the changes observed in F1 animals likely would not persist in the F2 generation. However, direct studies to evaluate the potential for transgenerational impacts of CS exposure, and possible impacts in humans, are warranted.

Our study has important implications in characterizing the potential mechanisms that underlie the elevated health risks observed in offspring of men who smoked prior to conception. The findings reported here are likely also applicable to understanding the risks of other environmental exposures that induce oxidative stress such as air pollution and some chemical exposures. Paternal preconception exposures to a variety of pharmacologic agents and pollutants, including nicotine, THC, morphine, and benzo[a]pyrene affect offspring phenotype, and often confer neurobehavioral consequences (Dai et al., 2017, Hawkey et al., 2019, Vallaster et al., 2017, Pachenari et al., 2019, Zhang et al., 2019, Viluksela and Pohjanvirta, 2019, Levin et al., 2019, McCarthy et al., 2018). In some cases, the impacts are transmitted across multiple generations (Zhang et al., 2019, Viluksela and Pohjanvirta, 2019). In this study, the absence of correlation between DMRs in F0 sperm and F1 sperm suggests that the affects we observed are likely not transmitted beyond the first generation, but we cannot rule out an alternate mechanism for transgenerational inheritance, including noncoding RNAs and chromatin features. While additional studies are necessary to fully characterize the long-term impacts of CS exposure, the current study significantly expands our understanding of paternal CS exposure impacts on offspring, and identifies a mechanism underlying CS-induced epigenetic changes in sperm.

## SUPPLEMENTAL FIGURE LEGEND

**Figure S1. Descriptive statistics of sperm parameters, fertility measures and animal weights for F0 animals and F1 animals.**

A) F0 animals- Both WT and *Nrf2*^*-/-*^ mice that were exposed to CS weighed significantly less than non-exposed control animals. Sperm concentration and motility were not significantly impacted by CS exposure, sperm concentration was lower in *Nrf2*^*-/-*^ than WT males, and conception was significantly delayed in CS-exposed *Nrf2*^*-/-*^mice compared with unexposed *Nrf2*^*-/-*^mice.

B) F1 animals- Neither growth trajectories nor sperm parameters were different in F1 animals based on paternal CS exposure. Abbreviations: CN-control *Nrf-/-*, SN-smoke-exposed *Nrf-/-*, CWT-control wild type animals, SWT-smoke-exposed wild type animals.

**Figure S2. Summary of the recovery, persistence or emergence of differentially methylated loci (A and B) and properties of recovery regions (C, D and E)**

A) Quantitative data indicate that the number of differentially methylated loci in CS-exposed mice compared with age matched controls does not diminish following a recovery period of up to 171 days.

B) When considering only loci that lost methylation in the CS-exposed group, about one third of differentially methylated loci persisted for the entire recovery period (white), one third returned to baseline levels (blue) and one third emerged as differentially methylated following a recovery period (pink). Contrastingly, for loci that gained methylation as a result of CS exposure, a smaller fraction of differentially methylated CpGs persisted or recovered while nearly half of differentially methylated loci emerged during the recovery period.

C) Impact of the initial methylation status and direction of change on methylation recovery. The hypomethylated DMRs (<25% DNAme) in which methylation increased and the hypermethylated DMRs (> 75% DNAme) in which methylation decreased with smoke exposure were more likely to recover. Regions displaying methylation between 25% and 75% Regions of intermediate DNAme were less likely to recover across both groups. Both comparisons were to the 3 days post-CS group.

D) In every category of DMR (shared, recovery or new) variation diminished as CpG density of a region increased. The impact of CpG density on variation was particularly apparent for regions in which DNAme decreased as a result of CS exposure and later recovered to baseline.

E) DMRs that displayed increased DNAme in CS-exposed animals and later recovered were generally regions of lower CpG density, while DMRs that gained methylation and subsequently recovered were generally at regions of higher CpG density. Timepoint 2 = 103-day recovery group, and timepoint 3 = 171-day recovery group. Both comparisons were to the 3-day recovery group. Dynamics were similar to those observed at 28 days post CS (see figure 1D-F).

**Figure S3. Chromatin properties at F0 sperm DMRs and variation in F1 prefrontal cortex gene expression.**

A) Chromatin accessibility was not predictive of the propensity for DMRs to recover or be maintained after removal of CS exposure.

B-C) Localization of B) H3K4me3 and C) H3K27me3 was likewise not associated with DMR recovery or maintenance.

D) A high correlation in DMRs was observed between CS-exposed WT mice and *Nrf2*^-/-^ whether or not they were exposed to CS. No correlation was observed between DMRs identified in the current study compared with previously published DMRs associated with vinclozolin exposure (VD2) and protein restricted diet (PR).

E) Scatter plot of F1 prefrontal cortex variation in gene expression in CS-exposed versus control *Nrf*^*-/-*^ mice, demonstrating that paternal CS exposure did not further elevate the increased gene expression variation in *Nrf*^*-/-*^ offspring beyond the effects of associated with genotype alone (see Figure 4A).

F) Scatter plot of F1 prefrontal cortex gene expression variation in CS-exposed WT offspring versus control *Nrf*^*-/-*^ demonstrating that control *Nrf*^*-/-*^ offspring exhibit a similar degree of variation as CS-exposed WT animals.

**Figure S4. Scatter plots of single CpG and regional sperm DNAme for control and smoked NRF versus control and smoked WT.**

A) CS-associated DNAme at all CpGs (left) and regions (right) across the genome.

B) CS exposure and *Nrf*^*-/-*^ do not induce broad, high magnitude genome-wide changes in sperm DNA methylation, and the number and magnitude of change for differentially methylated regions is similar across for CS-associated and *Nrf*^*-/-*^*-*associated DMRs.

## MATERIALS AND METHODS

### Animals

#### Animal care

All animal experiments were performed under protocols that were approved by the University of Utah Institutional Animal Care and Use Committee (protocol # 14-11006). All animals were obtained from Jackson Laboratories (Bar Harbor, ME)

#### Experimental Design

Six to seven-week-old mice were assigned to one of two groups: CS-exposed mice and non-exposed control mice. Following 60 days of CS exposure, mice were bred to unexposed CAST/EiJ female mice. Groups of animals were euthanized and tissues collected 3, 28, 103, and 171 days after removal from CS exposure. Sperm DNA methylation analysis was performed by RRBS on F0 exposed and control animals. Offspring derived from exposed and control males were euthanized at 14-17 weeks of age, and tissues were collected. DNA methylation analysis was performed on sperm and prefrontal cortex to investigate the impact of paternal smoking status on methylation patterns in offspring. In addition, RNAseq was performed on prefrontal cortex tissue to investigate the association between paternal smoking and neural gene expression.

#### Smoke exposure

All CS-exposed and control mice were age matched and smoking was initiated between 6 and 7 weeks of age. Mice were exposed to CS using a Teague Model TE-10 (Teague Enterprises, Woodland, CA) smoking machine, which produces a combination of side-stream and mainstream CS. A pump on the machine “puffs” each 3R4F University of Kentucky research cigarette for 2 seconds for a total of 9 puffs before ejection. The 2.5-hour daily exposure occurred for 5 consecutive days per week over a period of 60 days. The smoking chamber atmosphere was periodically sampled to confirm total particulate matter concentrations of approximately 150 mg/m^3^, the human equivalent of smoking approximately 10-20 cigarettes per day (Barrett et al., 2002).

#### Smoking and recovery experiments

To characterize the impact of smoke exposure on the sperm DNA methylome, and the capacity for smoke-induced sperm DNAme alterations to recover following removal of the insult, we exposed 40 C57BL/6J (Jackson Labs Stock # 000664) to cigarette smoke for comparison against 10 age-matched, non-smoked controls. Ten CS-exposed mice and the 10 non-exposed controls were euthanized and tissues collected within three days of the CS exposure period. Subsequent “recovery” groups of 10 CS-exposed animals were euthanized 28, 103 and 171 days after the exposure period (corresponding to approximately 0.8, 3 and 5 spermatogenic cycles). In addition to experiments with WT animals, ten age-matched *Nrf2*^-/-^ mice on a C57BL/6J genetic background (Jackson Labs Stock # 017009) were exposed to the same doses of CS for the same time period, and ten age-matched unexposed *Nrf2*^-/-^ mice were utilized as controls.

#### Offspring transmission experiments

Founder mice for heritability experiments included WT C57BL/6J mice (Jackson Labs Stock # 000664) that were exposed and not exposed to CS (n = 10-12 per group). Approximately one week after the exposure period, exposed and control males were introduced to 6-week old CAST/EiJ female mice (Jackson Labs Stock # 000928), and pairs were kept together until F1 litters were born, or for 7 weeks without conceiving, whichever came first. The motivation for outcrossing males to CAST/EiJ females was to leverage polymorphic alleles to enable attributing reads to a specific parent, however due to the large average spacing of informative SNPs in the CAST strain and the short sequencing reads inherent in Illumina sequencing we were unable to classify the large majority of reads based on parent-of-origin. We therefore analyzed the data without regard to parent-of-origin. F1 litters were weaned at approximately 21 days of age, and pups were regularly weighed until they were euthanized. F1 animals were euthanized at 14-17 weeks of age, and heart, lung, liver, kidney, brain, testis and epididymal sperm were collected for molecular studies.

#### Animal phenotyping

Following epididymal sperm extraction, sperm count and motility were assessed in CS-exposed and control F0 animals as well as F1 offspring. In addition, time to conception and litter size were compared between F0 groups. F1 offspring were evaluated for growth trajectory. For statistical analysis of growth trajectories between groups, animal weights were plotted against age for all pups within a group (C57BL/6J or *Nrf2*^-/-^). Models to fit the data were tested, and a logarithmic model generally yielded the highest r^2^. Theoretical weights were calculated for each weight event based on the model generated, and differences between theoretical and actual weight were calculated. A mean of average differences within an individual across weight events was calculated for each animal, and unpaired student’s t-test was used to compare these differences between smoked and non-smoked animals within each group. Differences in animal weights and sperm parameters were evaluated using two-tailed Student’s t-test, and two-tailed Fisher’s Exact tests were used to evaluate weekly differences in conception between groups. P < 0.05 was considered significant.

### Molecular analyses

#### Sperm collection and DNA extraction

Sperm was collected from the cauda epididymis and vas deferens immediately after euthanasia by scoring the tissue along the length of the tubules with a 28-G needle and gently pressing the tissue to expel the sperm mass. Tissues were then placed in a center-well dish in equilibrated Quinn’s medium (CooperSurgical, Trumbull, CT) supplemented with FBS in a humidified CO_2_ incubator for one hour. Following the swim out period, sperm concentration and motility were assessed on a Makler chamber and sperm were snap frozen in liquid nitrogen. Samples were subsequently thawed and subjected to a stringent somatic cell lysis protocol to ensure a pure population of sperm. Briefly, samples were passed through a 40 µM filter to remove cell and tissue clumps followed by two 14 ml washes with ddH_2_O and incubation for at least 60 minutes in somatic cell lysis buffer (0.1% SDS, 0.5% Triton X in ddH_2_O) at 4° C. Following somatic cell lysis and visual confirmation of the absence of contaminating cells, sperm DNA was extracted using the Qiagen AllPrep Universal kit (Hilden, Germany). Samples in cell lysis buffer were passed through a 28-gauge syringe multiple times to disrupt sperm membranes and liberate nucleic acids prior to extraction.

#### Prefrontal cortex dissection and nucleic acid extraction

Following euthanasia of F1 males (n = 8 per group), left brain hemispheres were dissected and placed in PreAnalytiX PaxGene (Hombrechtikon, Switzerland) tissue stabilizer and after 24 hours, fixed in PaxGene fixative and stored at −80° C. Samples were subsequently thawed and prefrontal cortex dissected under a stereo microscope according to the method described by Chiu et al. (Chiu et al., 2007). Tissue was then disrupted using a microcentrifuge pestle, and RNA and DNA were extracted using the Qiagen AllPrep Universal kit according to manufacturer’s protocols.

#### RRBS library construction

Following DNA extraction, Bioo Scientific NEXTflex Bisulfite Library Prep Kit for Illumina Sequencing (PerkinElmer, Austin, TX) was used for library preparation. To maximize coverage, we employed two separate restriction digests with MspI and TaqαI. Following digestion, products were pooled, and Klenow Fragment was utilized to create 3’A overhangs. DNA was subsequently purified with Zymo DNA Clean and Concentrate Columns (Irvine, CA) followed by ligation of Methylated Illumina PE Adapters and Ampure purification with SPRI beads. Purified products were Sodium Bisulfite Converted using ZymoResearch EZ DNA Methylation Gold Kit, and libraries were amplified over 20 cycles using Platinum Taq DNA polymerase (ThermoFisher, Waltham, MA), followed by a final Ampure purification (Beckman Coulter, Indianapolis, IN) and confirmation of library size range on a 2% agarose gel. DNA was submitted to the Huntsman Cancer Institute High Throughput genomic core for sequencing on a Hi-Seq 2500 (Illumina, San Diego, CA) using 50 cycle-single read chemistry. Four to six samples were sequenced per lane for a minimum of 35-million reads per sample.

#### RNAseq library construction

RNA extracted from F1 frontal cortices (n = 8 per group) was subjected to Illumina TruSeq Stranded RNA kit with Ribo-Zero Gold library preparation and subsequently sequenced on a Hi-Seq 2500 using 50 Cycle-Single Read Sequencing v4. Eight samples were sequenced per lane for a minimum of 25-million reads per sample.

#### Bioinformatics analyses

For genome wide DNA methylation analysis, sequence data from RRBS libraries was aligned to the mouse mm10 genome using the Bismark pipeline with special attention to RRBS specific issues, as noted in the Bismark User Guide and the Bismark RRBS Guide. Only CpGs where read coverage was greater than 8 for at least 4 biological replicates were considered “scoreable” for downstream analysis. Only CpGs with more than 5% change in methylation relative to control samples were classified as differentially methylated. When considering DMRs, only regions greater than 50 base-pairs in length with 3 or more scorable CpGs were analyzed. Then, one third of the CpGs within each analyzed region needed to be differentially methylated in order for a given region to be under consideration as a DMR. Finally, qualifying regions were classified as bonafide DMRs if there was more than 5% change in methylation relative to control samples. For genome wide gene expression analysis, sequencing data from RNASeq libraries was aligned using Novoalign. Aligned splice junction were converted to genomic coordinates and low quality and non-unique reads were further parsed using SamTranscriptomeParser (USeq; v8.8.8) under default settings. Stranded differential expression analysis was calculated with the USeq program DefinedRegionDifferentialSeq, which utilizes DESeq2 and the reference mm10. Normalized read count tables were then analyzed in R, along with all DNA methylation data. Integration and parsing of bed files or tables was performed in R. Generation of all figures and statistical analyses was accomplished using standard methods in R, with the exception of aggregate histone modification profiles, which were generated using Deeptools. Gene ontology analysis was performed using DAVID Functional Annotation Bioinformatics Resources. GSEA was performed on normalized gene expression read count tables using software development at the Broad Institute (http://software.broadinstitute.org/gsea/index.jsp).

## Data Access

Genomics data is available through the NIH GEO Datasets under accession number GSE133742

## Acknowledgements

We thank Dr. Kristin Murphy for thoughtful comments and critique during preparation of the manuscript.

## REFERENCES

Barber, R., Plumb, M. A., Boulton, E., Roux, I. & Dubrova, Y. E. 2002. Elevated mutation rates in the germ line of first- and second-generation offspring of irradiated male mice. Proceedings of the National Academy of Sciences of the United States of America, 99, 6877–82.

Barber, R. C. & Dubrova, Y. E. 2006. The offspring of irradiated parents, are they stable? Mutation research, 598, 50–60.

Barrett, E. G., Wilder, J. A., March, T. H., Espindola, T. & Bice, D. E. 2002. Cigarette smoke-induced airway hyperresponsiveness is not dependent on elevated immunoglobulin and eosinophilic inflammation in a mouse model of allergic airway disease. Am J Respir Crit Care Med, 165, 1410–8.

Behl, M., Rao, D., Aagaard, K., Davidson, T. L., Levin, E. D., Slotkin, T. A., Srinivasan, S., Wallinga, D., White, M. F., Walker, V. R., Thayer, K. A. & Holloway, A. C. 2012. Evaluation of the Association between Maternal Smoking, Childhood Obesity, and Metabolic Disorders: A National Toxicology Program Workshop Review. Environmental health perspectives.

Bosse, Y., Postma, D. S., Sin, D. D., Lamontagne, M., Couture, C., Gaudreault, N., Joubert, P., Wong, V., Elliott, M., Van Den Berge, M., Brandsma, C. A., Tribouley, C., Malkov, V., Tsou, J. A., Opiteck, G. J., Hogg, J. C., Sandford, A. J., Timens, W., Pare, P. D. & Laviolette, M. 2012. Molecular signature of smoking in human lung tissues. Cancer research, 72, 3753–63.

Breton, C. V., Byun, H. M., Wenten, M., Pan, F., Yang, A. & Gilliland, F. D. 2009. Prenatal tobacco smoke exposure affects global and gene-specific DNA methylation. American journal of respiratory and critical care medicine, 180, 462–7.

Brieno-Enriquez, M. A., Garcia-Lopez, J., Cardenas, D. B., Guibert, S., Cleroux, E., Ded, L., Hourcade Jde, D., Peknicova, J., Weber, M. & Del Mazo, J. 2015. Exposure to endocrine disruptor induces transgenerational epigenetic deregulation of microRNAs in primordial germ cells. PLoS One, 10, e0124296.

Busche, S., Shao, X., Caron, M., Kwan, T., Allum, F., Cheung, W. A., Ge, B., Westfall, S., Simon, M. M., Multiple Tissue Human Expression, R., Barrett, A., Bell, J. T., Mccarthy, M. I., Deloukas, P., Blanchette, M., Bourque, G., Spector, T. D., Lathrop, M., Pastinen, T. & Grundberg, E. 2015. Population whole-genome bisulfite sequencing across two tissues highlights the environment as the principal source of human methylome variation. Genome Biol, 16, 290.

Chiu, K., Lau, W. M., Lau, H. T., So, K. F. & Chang, R. C. 2007. Micro-dissection of rat brain for RNA or protein extraction from specific brain region. J Vis Exp, 269.

Cupul-Uicab, L. A., Skjaerven, R., Haug, K., Melve, K. K., Engel, S. M. & Longnecker, M. P. 2012. In utero exposure to maternal tobacco smoke and subsequent obesity, hypertension, and gestational diabetes among women in the MoBa cohort. Environmental health perspectives, 120, 355–60.

Dai, J., Wang, Z., Xu, W., Zhang, M., Zhu, Z., Zhao, X., Zhang, D., Nie, D., Wang, L. & Qiao, Z. 2017. Paternal nicotine exposure defines different behavior in subsequent generation via hyper-methylation of mmu-miR-15b. Sci Rep, 7, 7286.

Difranza, J. R., Aligne, C. A. & Weitzman, M. 2004. Prenatal and postnatal environmental tobacco smoke exposure and children’s health. Pediatrics, 113, 1007–15.

Dubrova, Y. E. 2003. Radiation-induced transgenerational instability. Oncogene, 22, 7087–93.

Dubrova, Y. E., Hickenbotham, P., Glen, C. D., Monger, K., Wong, H. P. & Barber, R. C. 2008. Paternal exposure to ethylnitrosourea results in transgenerational genomic instability in mice. Environmental and molecular mutagenesis, 49, 308–11.

Ford, E. S., Giles, W. H. & Dietz, W. H. 2002. Prevalence of the metabolic syndrome among US adults: findings from the third National Health and Nutrition Examination Survey. JAMA: the journal of the American Medical Association, 287, 356–9.

Ford, E. S., Giles, W. H. & Mokdad, A. H. 2004. Increasing prevalence of the metabolic syndrome among u.s. Adults. Diabetes Care, 27, 2444–9.

Fuentes, A., Munoz, A., Barnhart, K., Arguello, B., Diaz, M. & Pommer, R. 2010. Recent cigarette smoking and assisted reproductive technologies outcome. Fertility and sterility, 93, 89–95.

Glen, C. D. & Dubrova, Y. E. 2012. Exposure to anticancer drugs can result in transgenerational genomic instability in mice. Proceedings of the National Academy of Sciences of the United States of America, 109, 2984–8.

Hawkey, A. B., White, H., Pippen, E., Greengrove, E., Rezvani, A. H., Murphy, S. K. & Levin, E. D. 2019. Paternal nicotine exposure in rats produces long-lasting neurobehavioral effects in the offspring. Neurotoxicol Teratol, 74, 106808.

Holland, M. L., Lowe, R., Caton, P. W., Gemma, C., Carbajosa, G., Danson, A. F., Carpenter, A. A., Loche, E., Ozanne, S. E. & Rakyan, V. K. 2016. Early-life nutrition modulates the epigenetic state of specific rDNA genetic variants in mice. Science, 353, 495–8.

Ishii, T., Itoh, K., Takahashi, S., Sato, H., Yanagawa, T., Katoh, Y., Bannai, S. & Yamamoto, M. 2000. Transcription factor Nrf2 coordinately regulates a group of oxidative stress-inducible genes in macrophages. J Biol Chem, 275, 16023–9.

Jenkins, T. G., James, E. R., Alonso, D. F., Hoidal, J. R., Murphy, P. J., Hotaling, J. M., Cairns, B. R., Carrell, D. T. & Aston, K. I. 2017. Cigarette smoking significantly alters sperm DNA methylation patterns. Andrology.

Joubert, B. R., Haberg, S. E., Nilsen, R. M., Wang, X., Vollset, S. E., Murphy, S. K., Huang, Z., Hoyo, C., Midttun, O., Cupul-Uicab, L. A., Ueland, P. M., Wu, M. C., Nystad, W., Bell, D. A., Peddada, S. D. & London, S. J. 2012. 450K epigenome-wide scan identifies differential DNA methylation in newborns related to maternal smoking during pregnancy. Environmental health perspectives, 120, 1425–31.

Jung, Y. H., Sauria, M. E. G., Lyu, X., Cheema, M. S., Ausio, J., Taylor, J. & Corces, V. G. 2017. Chromatin States in Mouse Sperm Correlate with Embryonic and Adult Regulatory Landscapes. Cell Rep, 18, 1366–1382.

Kiziler, A. R., Aydemir, B., Onaran, I., Alici, B., Ozkara, H., Gulyasar, T. & Akyolcu, M. C. 2007. High levels of cadmium and lead in seminal fluid and blood of smoking men are associated with high oxidative stress and damage in infertile subjects. Biological trace element research, 120, 82–91.

Kohli, A., Garcia, M. A., Miller, R. L., Maher, C., Humblet, O., Hammond, S. K. & Nadeau, K. 2012. Secondhand smoke in combination with ambient air pollution exposure is associated with increasedx CpG methylation and decreased expression of IFN-gamma in T effector cells and Foxp3 in T regulatory cells in children. Clinical epigenetics, 4, 17.

Kulikauskas, V., Blaustein, D. & Ablin, R. J. 1985. Cigarette smoking and its possible effects on sperm. Fertility and sterility, 44, 526–8.

Laubenthal, J., Zlobinskaya, O., Poterlowicz, K., Baumgartner, A., Gdula, M. R., Fthenou, E., Keramarou, M., Hepworth, S. J., Kleinjans, J. C., Van Schooten, F. J., Brunborg, G., Godschalk, R. W., Schmid, T. E. & Anderson, D. 2012. Cigarette smoke-induced transgenerational alterations in genome stability in cord blood of human F1 offspring. FASEB journal: official publication of the Federation of American Societies for Experimental Biology, 26, 3946–56.

Levin, E. D., Hawkey, A. B., Hall, B. J., Cauley, M., Slade, S., Yazdani, E., Kenou, B., White, H., Wells, C., Rezvani, A. H. & Murphy, S. K. 2019. Paternal THC exposure in rats causes long-lasting neurobehavioral effects in the offspring. Neurotoxicol Teratol, 74, 106806.

Marchetti, F., Rowan-Carroll, A., Williams, A., Polyzos, A., Berndt-Weis, M. L. & Yauk, C. L. 2011. Sidestream tobacco smoke is a male germ cell mutagen. Proceedings of the National Academy of Sciences of the United States of America, 108, 12811–4.

Mccarthy, D. M., Morgan, T. J., Jr., Lowe, S. E., Williamson, M. J., Spencer, T. J., Biederman, J. & Bhide, P. G. 2018. Nicotine exposure of male mice produces behavioral impairment in multiple generations of descendants. PLoS Biol, 16, e2006497.

Moritsugu, K. P. 2007. The 2006 Report of the Surgeon General: the health consequences of involuntary exposure to tobacco smoke. American journal of preventive medicine, 32, 542–3.

Oberg, M., Jaakkola, M. S., Woodward, A., Peruga, A. & Pruss-Ustun, A. 2011. Worldwide burden of disease from exposure to second-hand smoke: a retrospective analysis of data from 192 countries. Lancet, 377, 139–46.

Pachenari, N., Azizi, H. & Semnaniann, S. 2019. Adolescent Morphine Exposure in Male Rats Alters the Electrophysiological Properties of Locus Coeruleus Neurons of the Male Offspring. Neuroscience, 410, 108–117.

Perera, F. & Herbstman, J. 2011. Prenatal environmental exposures, epigenetics, and disease. Reproductive toxicology, 31, 363–73.

Polyzos, A., Schmid, T. E., Pina-Guzman, B., Quintanilla-Vega, B. & Marchetti, F. 2009. Differential sensitivity of male germ cells to mainstream and sidestream tobacco smoke in the mouse. Toxicology and applied pharmacology, 237, 298–305.

Portela, A. & Esteller, M. 2010. Epigenetic modifications and human disease. Nat Biotechnol, 28, 1057–68.

Priskorn, L., Holmboe, S. A., Jacobsen, R., Jensen, T. K., Lassen, T. H. & Skakkebaek, N. E. 2012. Increasing trends in childlessness in recent birth cohorts - a registry-based study of the total Danish male population born from 1945 to 1980. International journal of andrology, 35, 449–55.

Rolland, M., Le Moal, J., Wagner, V., Royere, D. & De Mouzon, J. 2012. Decline in semen concentration and morphology in a sample of 26 609 men close to general population between 1989 and 2005 in France. Human reproduction.

Secretan, B., Straif, K., Baan, R., Grosse, Y., El Ghissassi, F., Bouvard, V., Benbrahim-Tallaa, L., Guha, N., Freeman, C., Galichet, L. & Cogliano, V. 2009. A review of human carcinogens--Part E: tobacco, areca nut, alcohol, coal smoke, and salted fish. The lancet oncology, 10, 1033–4.

Swan, S. H., Elkin, E. P. & Fenster, L. 1997. Have sperm densities declined? A reanalysis of global trend data. Environmental health perspectives, 105, 1228–32.

Tsaprouni, L. G., Yang, T. P., Bell, J., Dick, K. J., Kanoni, S., Nisbet, J., Vinuela, A., Grundberg, E., Nelson, C. P., Meduri, E., Buil, A., Cambien, F., Hengstenberg, C., Erdmann, J., Schunkert, H., Goodall, A. H., Ouwehand, W. H., Dermitzakis, E., Spector, T. D., Samani, N. J. & Deloukas, P. 2014. Cigarette smoking reduces DNA methylation levels at multiple genomic loci but the effect is partially reversible upon cessation. Epigenetics, 9, 1382–96.

Vallaster, M. P., Kukreja, S., Bing, X. Y., Ngolab, J., Zhao-Shea, R., Gardner, P. D., Tapper, A. R. & Rando, O. J. 2017. Paternal nicotine exposure alters hepatic xenobiotic metabolism in offspring. Elife, 6.

Venugopal, R. & Jaiswal, A. K. 1998. Nrf2 and Nrf1 in association with Jun proteins regulate antioxidant response element-mediated expression and coordinated induction of genes encoding detoxifying enzymes. Oncogene, 17, 3145–56.

Viluksela, M. & Pohjanvirta, R. 2019. Multigenerational and Transgenerational Effects of Dioxins. Int J Mol Sci, 20.

Word, B., Lyn-Cook, L. E., Jr., Mwamba, B., Wang, H., Lyn-Cook, B. & Hammons, G. 2012. Cigarette Smoke Condensate Induces Differential Expression and Promoter Methylation Profiles of Critical Genes Involved in Lung Cancer in NL-20 Lung Cells In Vitro: Short-Term and Chronic Exposure. International journal of toxicology.

Zeilinger, S., Kuhnel, B., Klopp, N., Baurecht, H., Kleinschmidt, A., Gieger, C., Weidinger, S., Lattka, E., Adamski, J., Peters, A., Strauch, K., Waldenberger, M. & Illig, T. 2013. Tobacco smoking leads to extensive genome-wide changes in DNA methylation. PLoS One, 8, e63812.

Zhang, W., Yang, J., Lv, Y., Li, S. & Qiang, M. 2019. Paternal benzo[a]pyrene exposure alters the sperm DNA methylation levels of imprinting genes in F0 generation mice and their unexposed F1-2 male offspring. Chemosphere, 228, 586–594.

