## Supplemental Figures for "Oxidative stress underlies heritable impacts of paternal cigarette smoke exposure"

### Figure S1

## A

| | M/ml sperm | Total motility | Days to conception | # conceived w/in 1 week | # conceived w/in 2 weeks | # conceived w/in 3 weeks | # conceived w/in 4 weeks | # never conceived | Pre-smoke weight | Weight after 4 wks treatment | Weight after 8 wks treatment |
|---|---|---|---|---|---|---|---|---|---|---|---|
| **CN** | 48.4 | 58.7 | 6.5 | 9.0 | 9.0 | 10.0 | 10.0 | 2.0 | 20.1 | 24.9 | 27.7 |
| **SN** | 40.2 | 56.5 | 11.8 | 3.0 | 3.0 | 4.0 | 5.0 | 6.0 | 21.4 | 21.1 | 22.7 |
| **CN vs SN p val** | 0.110 | 0.526 | 0.204 | **0.039** | **0.039** | **0.036** | 0.089 | 0.193 | 0.247 | **0.001** | **<0.0001** |
| **CWT** | 55.2 | 55.4 | 5.8 | 6.0 | 8.0 | 8.0 | 8.0 | 4.0 | 21.6 | 25.8 | 28.6 |
| **SWT** | 51.8 | 54.8 | 5.4 | 6.0 | 6.0 | 7.0 | 7.0 | 5.0 | 21.0 | 21.1 | 22.4 |
| **CWT vs SWT p val** | 0.618 | 0.868 | 0.908 | 1.000 | 0.608 | 1.000 | 1.000 | 1.000 | **0.029** | **<0.0001** | **<0.0001** |
| **NRF** | 44.5 | 57.6 | 8.3 | 12.0 | 12.0 | 14.0 | 15.0 | 8.0 | 20.9 | 23.0 | 25.2 |
| **WT** | 53.5 | 55.1 | 5.6 | 12.0 | 14.0 | 15.0 | 15.0 | 9.0 | 21.3 | 23.2 | 25.2 |
| **NRF vs WT p val** | **0.039** | 0.306 | 0.261 | 1.000 | 0.773 | 1.000 | 1.000 | 1.000 | 0.444 | 0.757 | 0.975 |
| **Smoke** | 46.2 | 55.6 | 8.1 | 9.0 | 9.0 | 11.0 | 12.0 | 11.0 | 21.0 | 21.1 | 22.5 |
| **No Smoke** | 51.8 | 57.0 | 6.2 | 15.0 | 17.0 | 18.0 | 18.0 | 6.0 | 21.4 | 25.6 | 28.4 |
| **Smoke vs No Smoke p val** | 0.207 | 0.556 | 0.432 | 0.148 | **0.042** | 0.075 | 0.135 | 0.135 | 0.221 | **<0.0001** | **<0.0001** |

## B

| | Litter size | M/ml sperm | Total sperm motility | Postnatal weight (g) week 4 | Postnatal weight (g) week 6 | Postnatal weight (g) week 10 |
|---|---|---|---|---|---|---|
| **CN** | 5.3 | 25.7 | 36.1 | 11.6 | 16.4 | 17.4 |
| **SN** | 6.0 | 26.5 | 33.0 | 12.3 | 16.6 | 18.7 |
| **CN vs SN p val** | 0.379 | 0.851 | 0.409 | 0.351 | 0.727 | 0.100 |
| **CWT** | 5.0 | 31.3 | 27.3 | 12.1 | 16.2 | 17.4 |
| **SWT** | 5.2 | 33.6 | 34.0 | 12.7 | 16.8 | 17.8 |
| **CWT vs SWT p val** | 0.723 | 0.708 | 0.207 | 0.254 | 0.260 | 0.354 |
| **NRF** | 5.5 | 26.0 | 34.0 | 11.8 | 16.4 | 17.6 |
| **WT** | 5.1 | 32.5 | 30.5 | 12.5 | 16.4 | 17.5 |
| **NRF vs WT p val** | 0.356 | 0.063 | 0.264 | 0.194 | 0.979 | 0.828 |
| **Smoke** | 5.6 | 29.8 | 32.0 | 12.5 | 16.7 | 18.0 |
| **No Smoke** | 5.2 | 27.3 | 33.2 | 11.7 | 16.3 | 17.4 |
| **Smoke vs No Smoke p val** | 0.424 | 0.469 | 0.700 | 0.104 | 0.288 | 0.105 |

### Figure S2

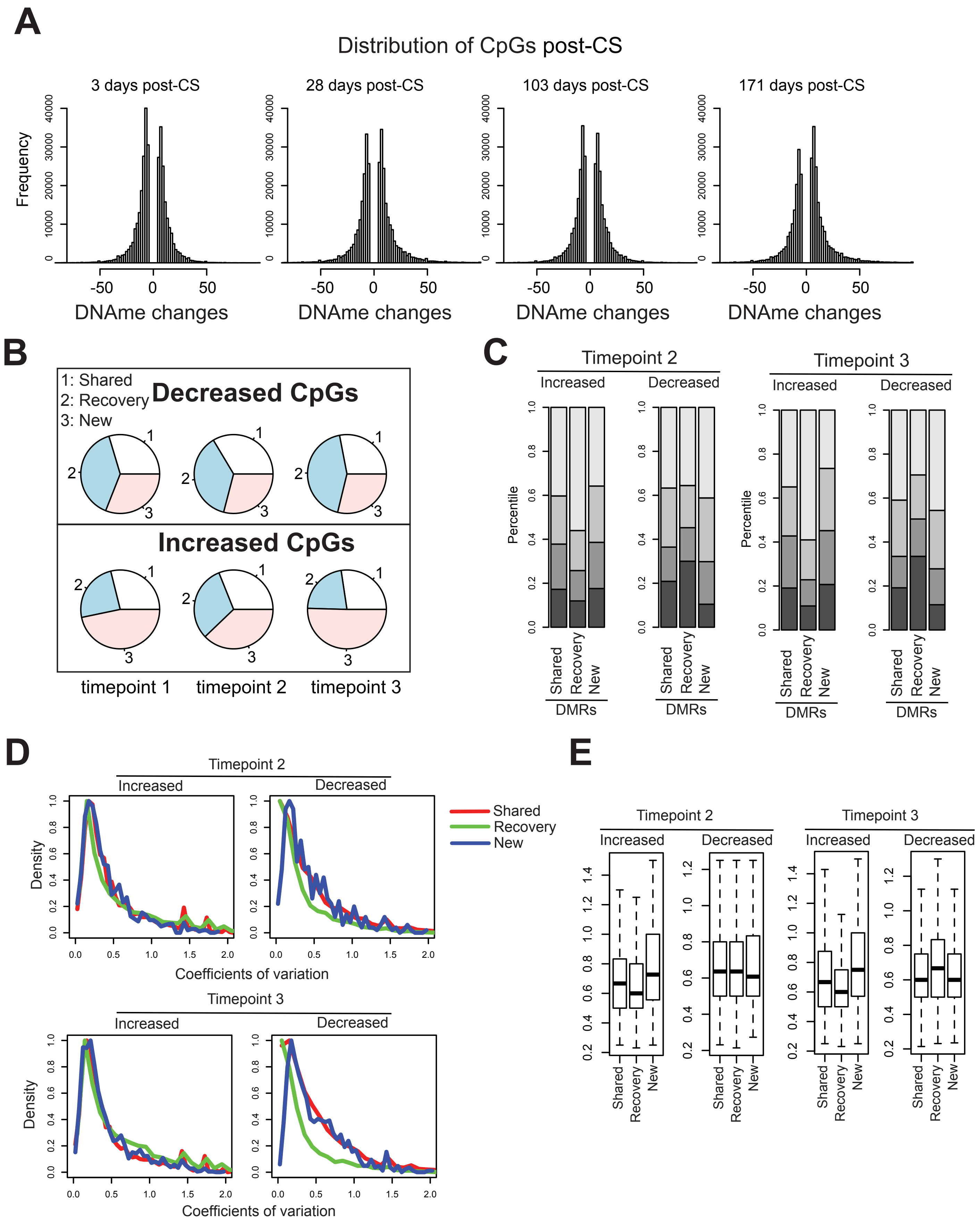
A
Distribution of CpGs post-CS
3 days post-CS
28 days post-CS
103 days post-CS
171 days post-CS
Frequency
DNAme changes
B
1: Shared
2: Recovery
3: New
Decreased CpGs
Increased CpGs
timepoint 1
timepoint 2
timepoint 3
C
Timepoint 2
Timepoint 3
Increased
Decreased
Percentile
Shared
Recovery
New
DMRs
D
Timepoint 2
Timepoint 3
Increased
Decreased
Density
Coefficients of variation
Shared
Recovery
New
E
Timepoint 2
Timepoint 3
Increased
Decreased
Shared
Recovery
New

### Figure S3

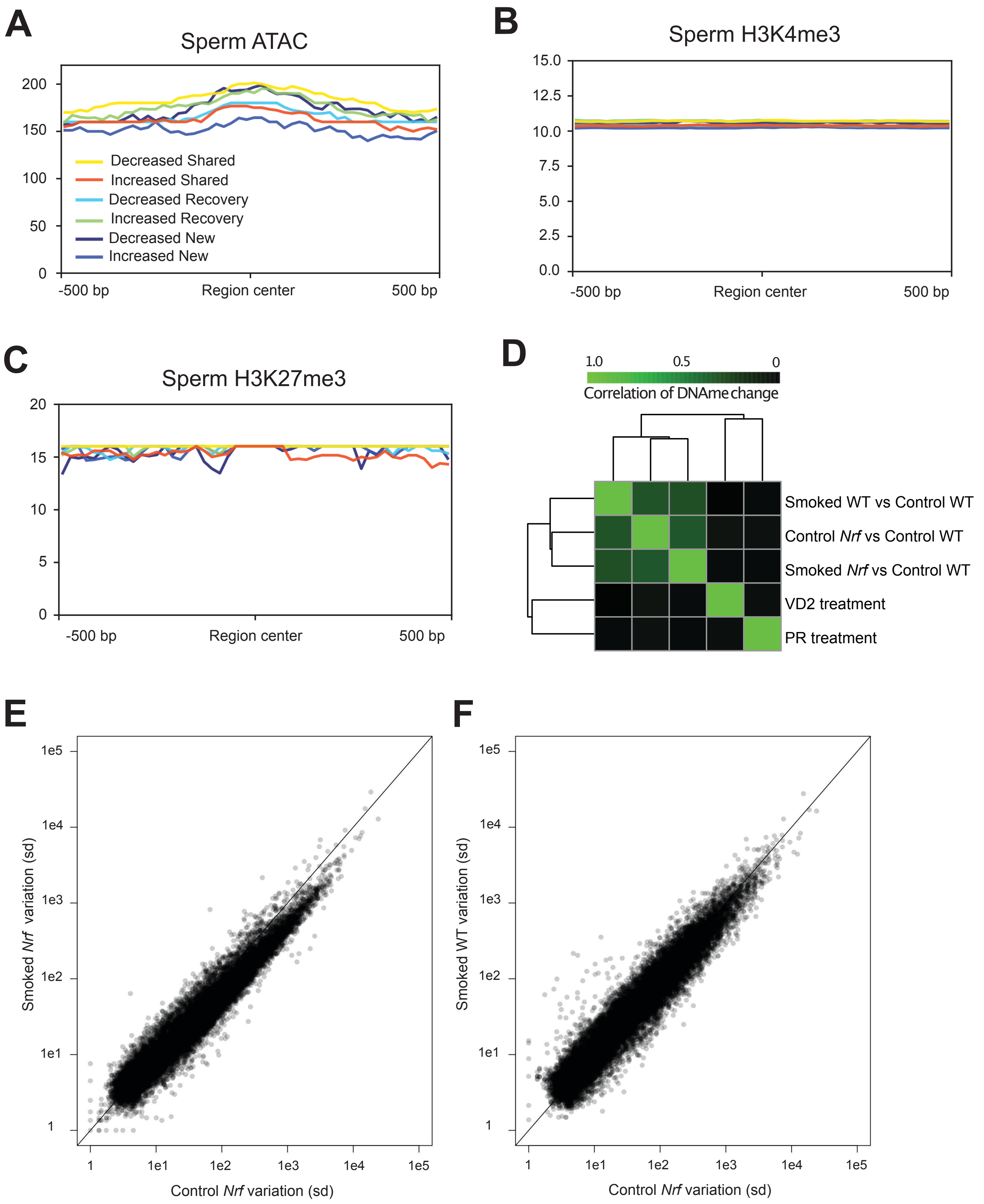
A
Sperm ATAC
200
150
100
50
0
Decreased Shared
Increased Shared
Decreased Recovery
Increased Recovery
Decreased New
Increased New
-500 bp
Region center
500 bp
B
Sperm H3K4me3
15.0
12.5
10.0
7.5
5.0
2.5
0.0
-500 bp
Region center
500 bp
C
Sperm H3K27me3
20
15
10
5
0
-500 bp
Region center
500 bp
D
1.0
0.5
0
Correlation of DNAme change
Smoked WT vs Control WT
Control Nrf vs Control WT
Smoked Nrf vs Control WT
VD2 treatment
PR treatment
E
Smoked Nrf variation (sd)
1e5
1e4
1e3
1e2
1e1
1
Control Nrf variation (sd)
F
Smoked WT variation (sd)
1e5
1e4
1e3
1e2
1e1
1
Control Nrf variation (sd)

Figure S4

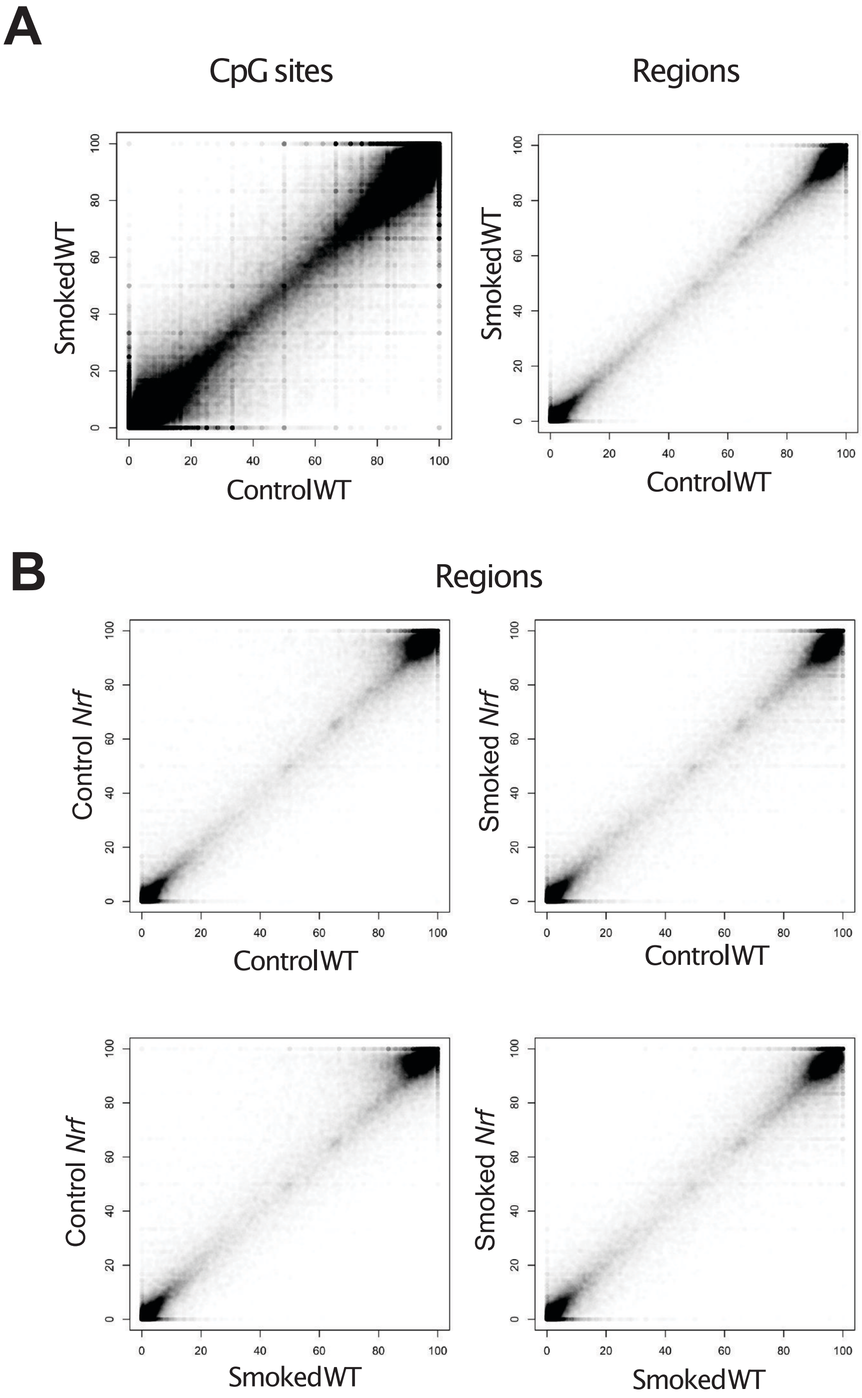

A
CpG sites
Regions
SmokedWT
ControlWT
B
Regions
Control *Nrf*
Smoked *Nrf*
ControlWT
SmokedWT
0
20
40
60
80
100
